# Progesterone regulates hypothalamic-pituitary-thyroid axis

**DOI:** 10.1101/166702

**Authors:** Chunyun Zhong, Kewen Xiong, Xin Wang

## Abstract

Progesterone is a natural steroid hormone excreted by animals and humans, which has been frequently detected in the aquatic ecosystems. The effects of the residual progesterone on fish are unclear. In this study, we aimed to examine the effects of progesterone on the hypothalamic-pituitary-thyroid (HPT) axis by detecting the gene transcriptional expression levels. Zebrafish embryos were treated with different concentrations of progesterone from 12 hours post-fertilization (hpf) to 120 hpf. Total mRNA was extracted and the transcriptional profiles of genes involved in HPT axis were examined using qPCR. The genes related to thyroid hormone metabolism and thyroid hormone synthesis were up-regulated in zebrafish exposed to progesterone. These results indicated that progesterone affected the mRNA expression of genes involved in the HPT axis, which might interrupt the endocrine system in zebrafish. Our data also suggested that zebrafish is a useful tool for evaluating the effects of chemicals on the thyroid endocrine system.

## Introduction

Natural and synthetic steroid hormones are active endocrine disrupters, which have been detected in the aquatic systems [1-5]. These endocrine disrupters have the potential effects on the fish reproductive system at very low concentration, such as environmental levels [6-9]. Progesterone is a steroid hormone that impairs the reproduction in animals and humans [10-12]. Previous studies report that progesterone can affect the meiotic oocyte maturation in human, mammals and female fish [13-18].

Low levels of progesterone and its metabolites, which are originally excreted by human and mammals, have been detected in the aquatic ecosystems including surface water and rivers [19-25].

Thyroid hormones play an essential role in the maintenance of tissues and biological functions in vertebrates [26-31]. In fish, the HPT axis regulates the thyroid endocrine system by modulating their homeostasis [32-35]. In this study, we investigated the effects of progesterone on the transcriptional profiles of genes involved in HPT axis of zebrafish. Our results confirmed that progesterone can cause disruption to the thyroid system by impairing the endocrine homeostasis.

## Materials and methods

### Chemicals

Progesterone was purchased from Sigma-Aldrich (Cat NO. P0130-25G) and dissolved in dimethyl sulfoxide to make the stock solution.

### Zebrafish maintenance and drug treatment

Wild type AB line zebrafish were raised and kept under standard laboratory conditions at 28 ± 0.5 °C. Adult fish were naturally crossed and normally developed embryos were collected. Embryos were randomly divided into each group containing different concentrations of progesterone (0, 1, 10, 100 and 1000 ng/L). The eggs were treated with progesterone from 12 hours post-fertilization (hpf) to 120 hpf. Water was changed twice per day and exposed-eggs were collected for mRNA analysis at 5 days post-fertilization (dpf). Control group was treated with 0.05% DMSO.

### Quantitative real-time PCR (qPCR)

Quantitative real-time PCR was performed as described somewhere else [36]. Fifty zebrafish larvae per group were collected and homogenized, total RNA was extracted using Direct-zol^TM^ RNA MiniPrep Kit (Zymo Research) following manufacturer’s protocol. First-strand cDNA was synthesized using SuperScript III First-Strand Synthesis System (Thermo Fisher Scientific). The qPCR was performed using SYBR Green Real-Time PCR Master Mixes (Thermo Fisher Scentific) and analyzed on a QuantStudio^TM^ 6 Flex Real-Time PCR System. The primer sequences of target genes were listed somewhere else. The amplification protocol was as follows: denaturation at 95 °C for 15 min, followed by 40 cycles of 95 °C for 10 s, 60 °C for 60 s.

### Statistical analysis

The normality and homogeneity of variance were tested using Kolmogorov-Smirnov and Levene’s tests, respectively. Differences between the control and drug-treated groups were evaluated by on-way analysis of variance (ANOVA), followed by Tukey’s test. P< 0.05 was considered statistically significant.

## Results

In order to investigate the effect of progesterone on the mRNA expression of genes involved in HPT axis of zebrafish, embryos were exposed to different concentrations of progesterone (0, 1, 10, 100 and 1000 ng/L), and mRNA expression profiles were examined at 5 dpf. Treated with 100 ng/L progesterone significantly increased the gene expression of sodium/iodide symporter (slc5a5) and thyroid-stimulating hormone beta (tshβ). The gene expression involved in the HPT axis had no significant changes in the 1 and 10 ng/L progesterone-treated groups. Treatment with higher concentrations (1000 ng/L) significantly induced the expression of NK2 homeobox 1a (nkx2.1), paired box protein 8 (pax8) and uridinediphosphate glucuronosyltransferase (ugt1ab).

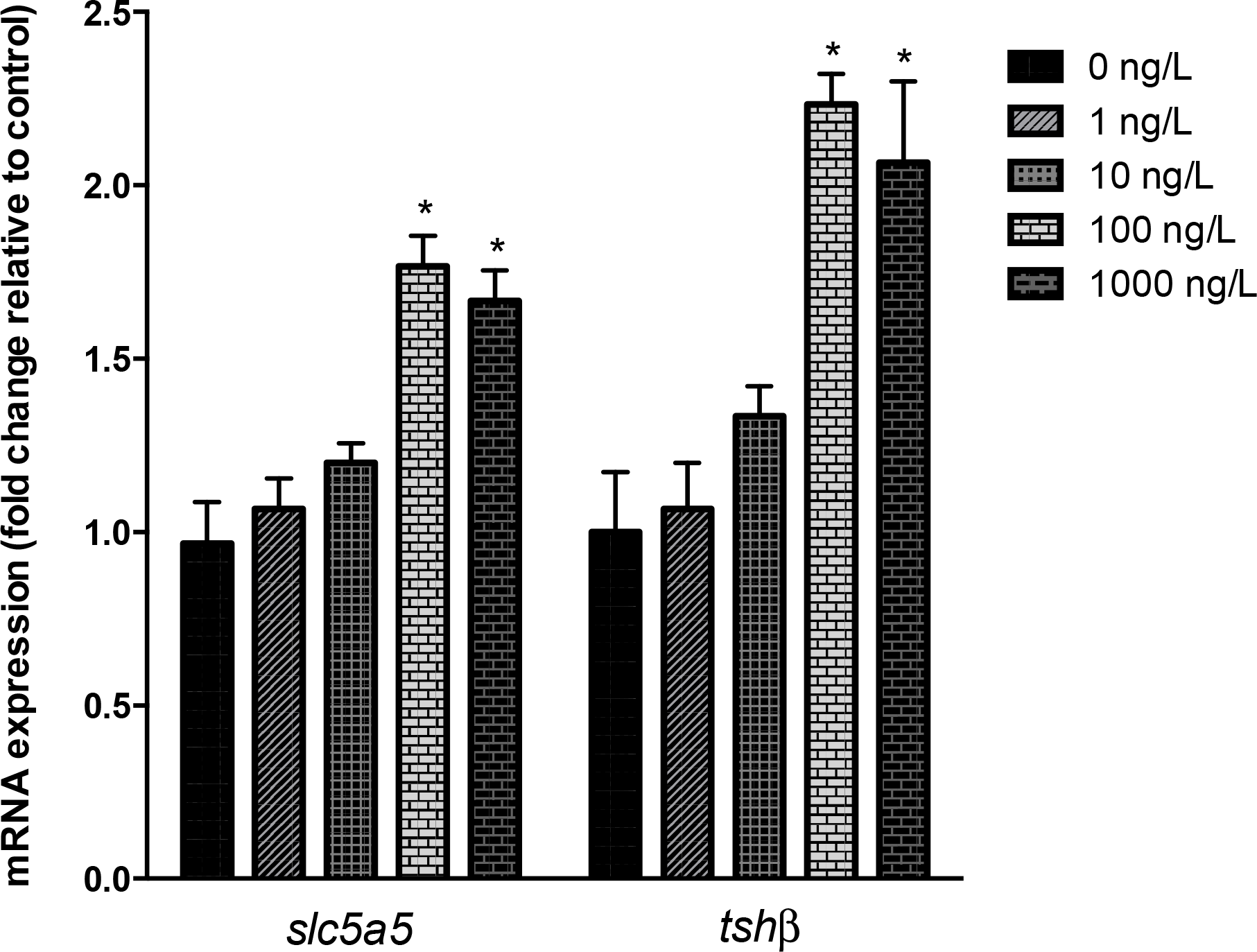

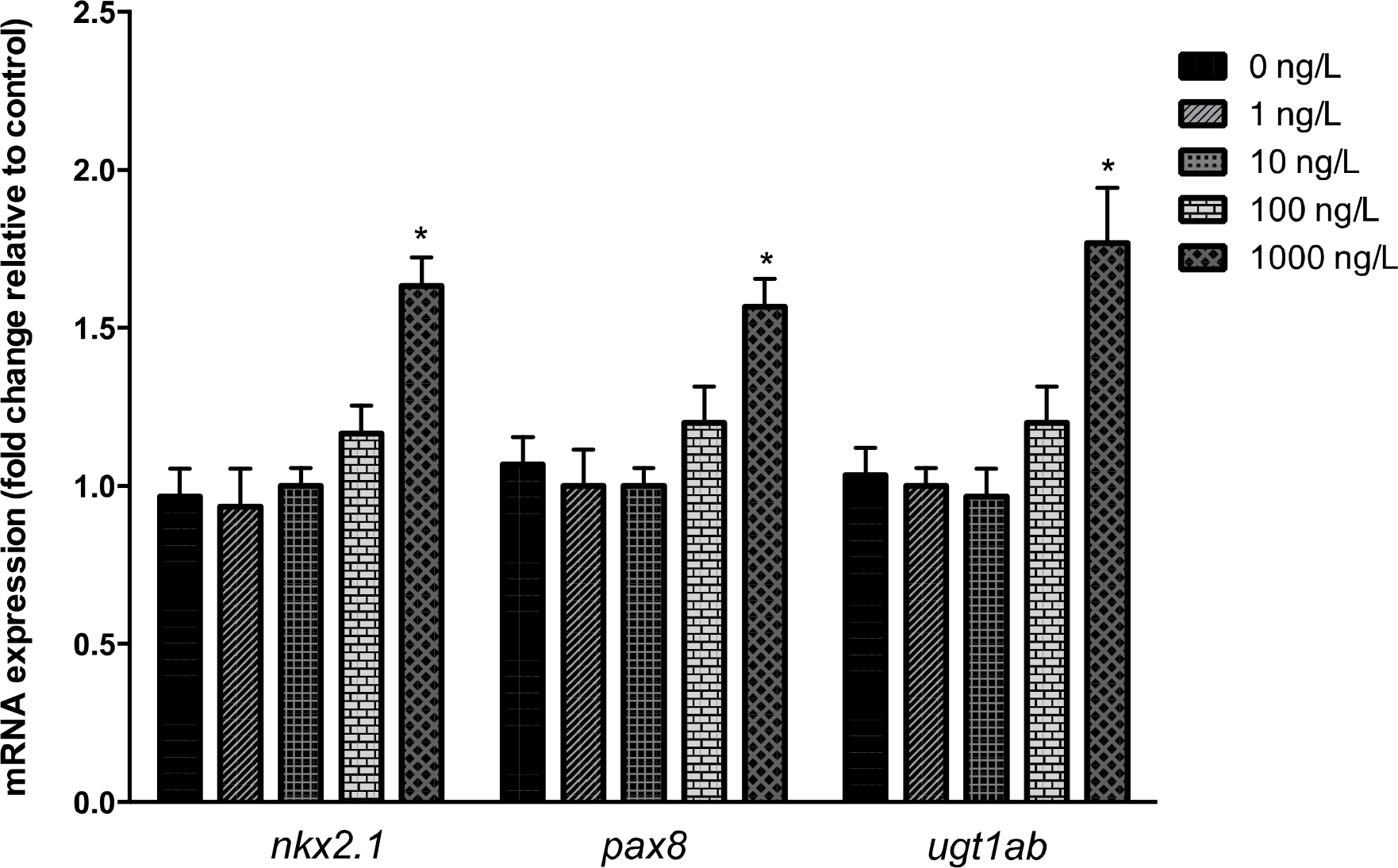

## Discussion

Very little is known of the effects of progesterone on the thyroid endocrine system in fish. In this study, zebrafish embryos were used to evaluate effects of progesterone on expression of genes involved in the HPT axis. Exposure to 100 ng/L progesterone significantly increased slc5a5 and tshβ expression. The gene slc5a5 and tshβ are involved in thyroid hormone synthesis pathways [37-39].

Previous studies have reported that tshβ is a useful biomarker for investigating the function of thyroid system [40, 41]. In this study, the tshβ mRNA expression was significantly increased when exposed to progesterone. Moreover, higher concentration of progesterone increased nkx2.1, pax8 and ugt1ab expression. Genes regulate the thyroid development (nkx2.1 and pax8) and thyroid hormone synthesis (slc5a5) were significantly upregulated after exposed to progesterone, these results suggested that HPT axis in the 5 dpf zebrafish is sensitive to chemical treatment, that can be used to examine the effects of chemicals on the thyroid endocrine system.

Taken together, treatment with progesterone changed the gene expression levels involved in the HPT axis, indicating an overt endocrine-disrupting activity. This study showed that 5 dpf zebrafish can be used to evaluate the effects of chemicals on the thyroid endocrine system.

